# Binding Interactions between RBD of Spike-Protein and Human ACE2 in Omicron variant

**DOI:** 10.1101/2022.02.10.480009

**Authors:** Bahaa Jawad, Puja Adhikari, Rudolf Podgornik, Wai-Yim Ching

**Author notes:** Corresponding Author: Wai-Yim Ching.

## Abstract

Emergence of new *SARS-CoV-2 Omicron VOC* (*OV*) has exacerbated the COVID-19 pandemic due to a large number of mutations in the spike-protein, particularly in the receptor-binding domain (RBD), resulting in highly contagious and/or vaccine-resistant strain. Herein, we present a systematic analysis based on detailed molecular dynamics (MD) simulations in order to understand how the OV RBD mutations affect the ACE2 binding. We show that the OV RBD binds to ACE2 more efficiently and tightly due predominantly to strong electrostatic interactions, thereby promoting increased infectivity and transmissibility compared to other strains. Some of OV RBD mutations are predicted to affect the antibody neutralization either through their role in the S-protein conformational changes, such as S371L, S373P, and S375F, or through changing its surface charge distribution, such as G339D, N440K, T478K, and E484A. Other mutations, such as K417N, G446S, and Y505H, decrease the ACE2 binding, whereas S447N, Q493R, G496S, Q498R, and N501Y tend to increase it.

**TOC GRAPHICS:** 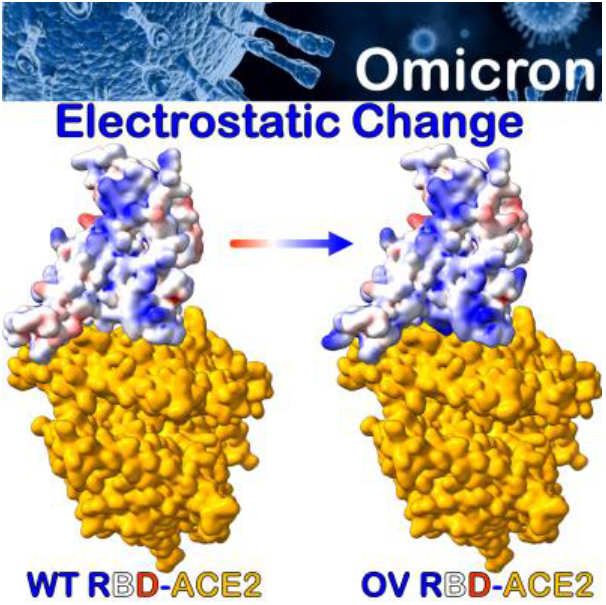

## 1. Introduction

Since the outbreak of COVID-19 pandemic its pathogen SARS-CoV-2 has continuously mutated and evolved, resulting in the emergence of major variants of concern (VOC). These VOC have been observed to alter the virus characteristics such as infectivity, transmissibility, antigenicity, and pathogenicity.^1^ The most recently identified SARS-CoV-2 VOC is the *Omicron variant (OV)* (B.1.1.529) which has quickly become the dominant strain.^2,3^ It has the highest number of amino acid (AA) mutations of any known SARS-CoV-2 VOC, with over 30 mutations in the spike (S) protein, of which 15 are in the receptor-binding domain (RBD),^4^ the main target for vaccine and treatment developments.^5–8^ The presence of many mutations in OV S-protein has raised concerns about elevated transmissibility, immunological escape, and vaccine and treatment failures.^9–15^ OV has been identified to contain several key mutations observed also in other SARS-CoV-2 VOC, such as K417N, E484A, and N501Y, that change the virus sensitivity to neutralization or increase the infectivity.^16^ Moreover, it contains many novel mutations that have not been observed previously, and their biological effects are largely unknown. Since the binding between RBD and the angiotensin-converting enzyme-2 (ACE2) facilitates viral entry, initiating the infection process, the fundamental understanding of how the OV RBD interacts with ACE2 is pivotal for understanding the viral infection mechanism and its evolution, as well as for therapeutic development of effective means to reduce its spread.

The present study aims to investigate how OV mutations affect the binding strength between RBD and ACE2 by highlighting the role of each mutation, its underlying mechanism and the pertinent binding driving forces. All-atom molecular dynamics (MD) simulations in explicit solvent have been implemented in order to study the dynamics and binding mechanism of the RBD-ACE2 system, followed by the molecular mechanics (MM) generalized Born surface area (MM-GBSA) method to predict the binding affinity and the binding profile. The results are compared with the previously reported analysis of the unmutated, wild type (WT) RBD-ACE2 system^17^, in order to assess the effect of each mutation and the nature of its interaction based on the per-residue and pairwise decomposition schemes.

## 2. Computational Modeling and Methods

We implement a procedure, previously developed for both Alpha, and Beta VOC, to build a computational model of the Omicron RBD-ACE2 system.^17^ We briefly summarize it as follows. *First*, the interface structure of RBD-ACE2 complex (PBD ID:6M0J) is used as a template to create the OV RBD-ACE2 system, including all 15 RBD mutations^18^: G339D, S371L, S373P, S375F, K417N, N440K, G446S, S477N, T478K, E484A, Q493R, G496S, Q498R, N501Y, and Y505H.^4^ The last 10 mutations are subsequently exposed to the ACE2 receptor as shown in **Figure**

To substitute the AAs of WT with mutated AAs of OV, we used the Dunbrack backbone-dependent rotamer library^19^ implemented by UCSF Chimera,^20^ which also allows for a careful control of backbone dihedral angles. For example, the torsion angles of K417N and N501Y mutations from the available RBD Beta VOC have been used in our OV model ^21^, as well as the angles of T478K mutation in the Delta variant.^22^ *Second*, the resulting OV RBD-ACE2 complex is solvated with 27000 explicit water molecules together with the appropriate number of ions (1 Zn^+2^, 1 Cl^-^, and 22 Na^+^) using the TIP3P explicit water model, implemented in AMBER^23,24^, with the most recent ff14SB force field used for parameterizations of intermolecular and intramolecular interactions in the OV complex.^25^ *Third*, the same MD steps of minimization, heating, equilibration, two MD production runs of over 100 ns were reproduced.^17^ *Finally*, the MM-GBSA method^17,26–29^ was applied to compute the *binding free energy* (BFE) and to quantify the complete binding profile^17^, allowing for a per-residue and pairwise BFE decompositions to identify the role and the nature of interaction for each mutation in the OV model.

## 3. Results and Discussion

**Figure 2(a)** shows the root-mean-square deviation (RMSD) for two replicate MD simulations of the OV, compared with the WT. All MD simulations have reached stability as shown by the stationary density distributions of RMSD and small standard deviations. The averaged RMSDs of OV complex, ACE2, RBD and RBM are 2.24 ± 0.22 Å, 1.98 ± 0.19 Å, 2.15 ± 0.2 Å, 1.63 ± 0.23 Å, respectively, while for WT they are 2.53 ± 0.29 Å, 2.32 ± 0.29 Å, 1.85 ± 0.12 Å, and 1.38 ± 0.12 Å, respectively.^17^ The average RMSDs of the 15 OV RBD mutations thus lead to a more stable interfacial complex with ACE2, even though the RMSDs of RBD and RBM themselves are relatively higher than the corresponding values of WT. In comparison with WT, the OV mutations are seen to only slightly change the RBD structure. This is even more pronounced when their RMSFs are compared, as shown in **Figure S1** in Supporting information (SI). The average RMSF of mutated RBD in OV is 1.45 Å vs 1.29 Å of WT RBD, indicating that the OV RBD is relatively less rigid than WT RBD, thus allowing for larger fluctuations. **Figure 2(b)** compares the RMSF of only the mutated AAs in OV vs WT. The average RMSF for RBM in both strains is calculated and found to be very similar (1.15 and 1.14 Å for OV and WT respectively). This demonstrates that the lesser rigidity of OV RBD is due to RBD core mutations rather than RBM mutations. In particular, D339, L371, P373, and F375 are relatively more flexible than their WT counterparts.

**Figure 1.**
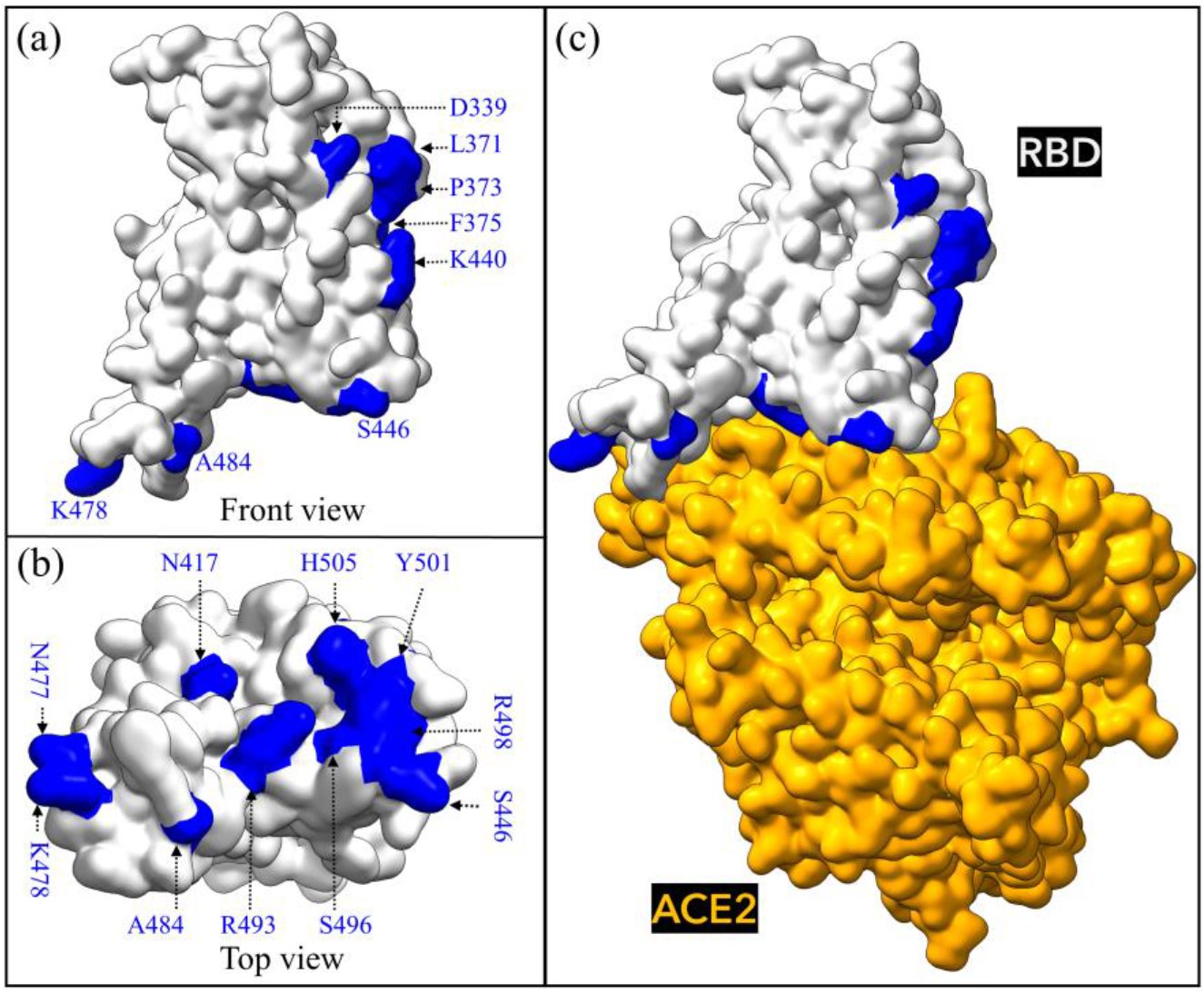
Omicron RBD-ACE2 interface system. (a) Front view of unbound Omicron variant (OV) RBD with all 15 mutations shown in blue and (b) Top view. (c) Bound OV RBD-ACE2 model.

**Figure 2.**
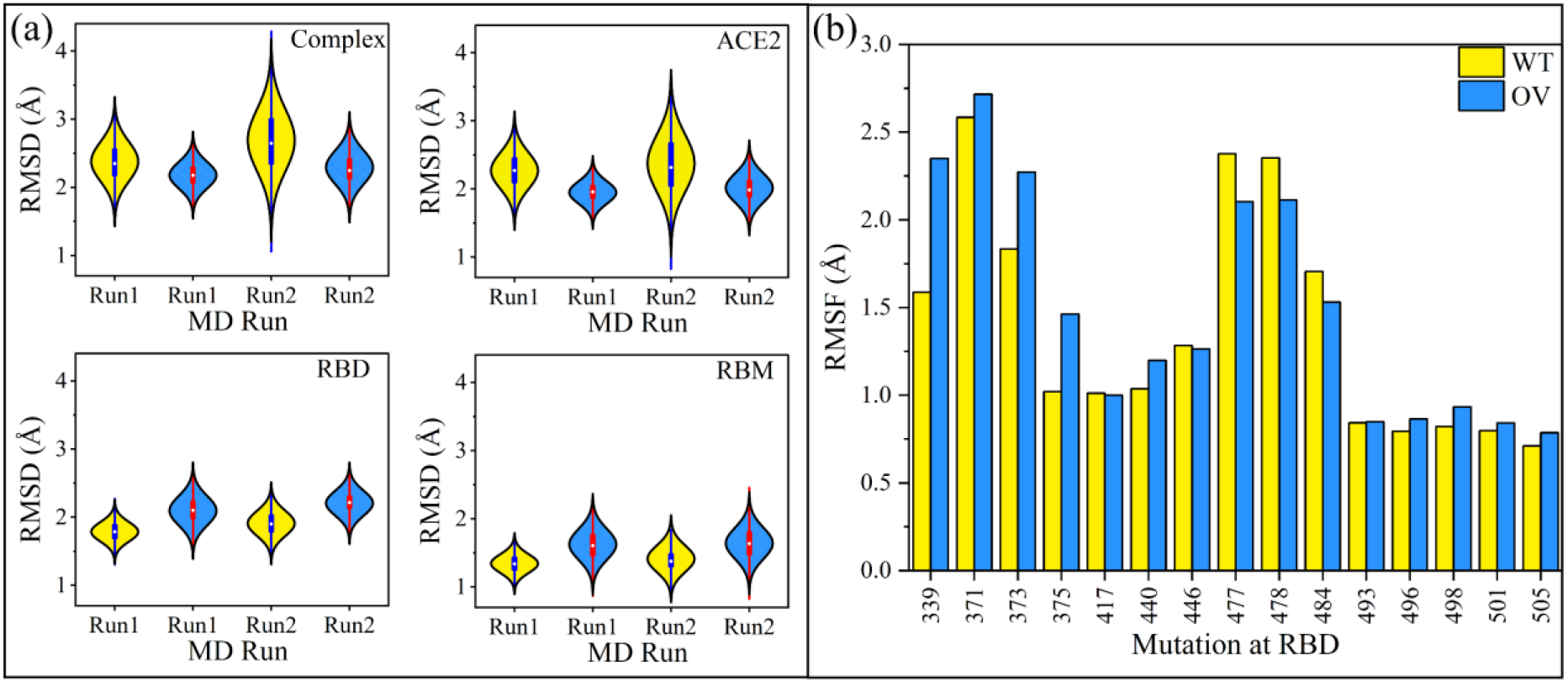
Stability of the OV RBD-ACE2 complex and slight conformation changes of mutated residues at RBD core. (a) RMSD violin plots of complex, ACE2, RBD and RBM from OV and WT for both MD runs. (b) RMSF for 15 residues that are mutated in OV and compared with WT.

This is consistent with a recent study based on the cryo-EM S-protein structure, showing that the L371, P373, and F375 significantly alter the conformation and mobility of RBD^12^, thus demonstrating that the computational MD simulations can confirm and reproduce what is observed experimentally.

### 3.1 Omicron RBD binds ACE2 stronger than WT RBD

The MM-GBSA method has been applied to calculate BFE of OV RBD-ACE2 system at 310 K (37 °C), with neutral pH (7.4) and 0.15 M uni-univalent NaCl salt concentration. **Table 1** presents the BFE and its decomposition for OV, and compared with WT.^17^ It shows that the OV RBD binds ACE2 more strongly than WT, with relative binding energy of −1.3 kcal/mol, consistent with recent experimental and computational studies.^11–13,30–32^ Interestingly, our predicted BFE value of −14.16 kcal/mol for OV falls between the Beta and Alpha BFE values of −13.5 and −14.7 kcal/mol, respectively ^17^ The complete thermodynamic decomposition listed in **Table 1** shows the Coulomb electrostatic interaction (ΔE_ele_) of OV to be more than twice that of WT. The increase in ΔE_ele_ of OV is mainly the result of five AAs at the RBM changing from polar to positively charged residues (N440K, T478K, Q493R, Q498R, and Y505H). The electrostatic component is thus seen as the main reason behind the more effective binding of ACE2 and OV RBD, which may also elucidate the reason behind the highly contagious nature of OV (**Figure S2**). ΔE_ele_ of OV, however, creates a higher desolvation energy (ΔG_GB_) that is indispensable in the formation process and cannot be avoided (**Table 1**). On the other hand, the van der Waals interaction (ΔEvdW) plays a key role in stabilizing and governing the RBD-ACE2 interaction as well as enhancing their binding by gaining −2.57 kcal/mol as compared to WT.

**Table 1.** Decomposition of BFE (kcal.mol-1) of RBD-ACE2 complex in OV and WT.

| Energy | OV (SEM) | WT (SEM)<sup>1</sup> | $\Delta\Delta G^2$ |
| --- | --- | --- | --- |
| $\Delta E_{\text{vdw}}$ | -92.67 (0.1) | -90.10 (0.1) | -2.57 |
| $\Delta E_{\text{ele}}$ | -1480.65 (0.69) | -700.92 (0.6) | -779.73 |
| $\Delta E_{\text{MM}}$ | -1573.31 (0.72) | -791.03 (0.7) | -782.28 |
| $\Delta G_{\text{GB}}$ | 1529.00 (0.69) | 748.49 (0.6) | 780.51 |
| $\Delta G_{\text{SA}}$ | -13.18 (0) | -13.21 (0) | 0.03 |
| $\Delta G_{\text{sol}}$ | 1515.82 (0.69) | 735.28 (0.6) | 780.54 |
| -T$\Delta S$ | 43.33 | 42.89 | 0.44 |
| $\Delta G_{\text{bind}}$ | -14.16 (0.1) | -12.86 (0.1) | -1.30 |

<sup>1</sup>All values are taken from ref.17 and SEM is the standard error of the mean. <sup>2</sup> $\Delta\Delta G = \Delta G_{\text{OV}} - \Delta G_{\text{WT}}$ .

### 3.2 Role and nature of interaction due to mutations in Omicron RBD

To gain a better understanding of the nature and impact of each mutation on the BFE in RBD-ACE2, in terms of per-residue fractions, decomposition schemes have been implemented and are shown in **Figure S3** for all OV and WT single AAs and AA-AA pairs. **Figure 3(a)** shows the per-residue BFE decomposition change for the 15 mutations between OV and WT (ΔΔG = ΔG_OV_ - ΔG_WT_), while their AA-AA interaction pair maps are displayed in **Figures 3(b)**. These mutations are divided into three groups based on their influence on binding: neutral, decreased and increased binding.

**Figure 3.**
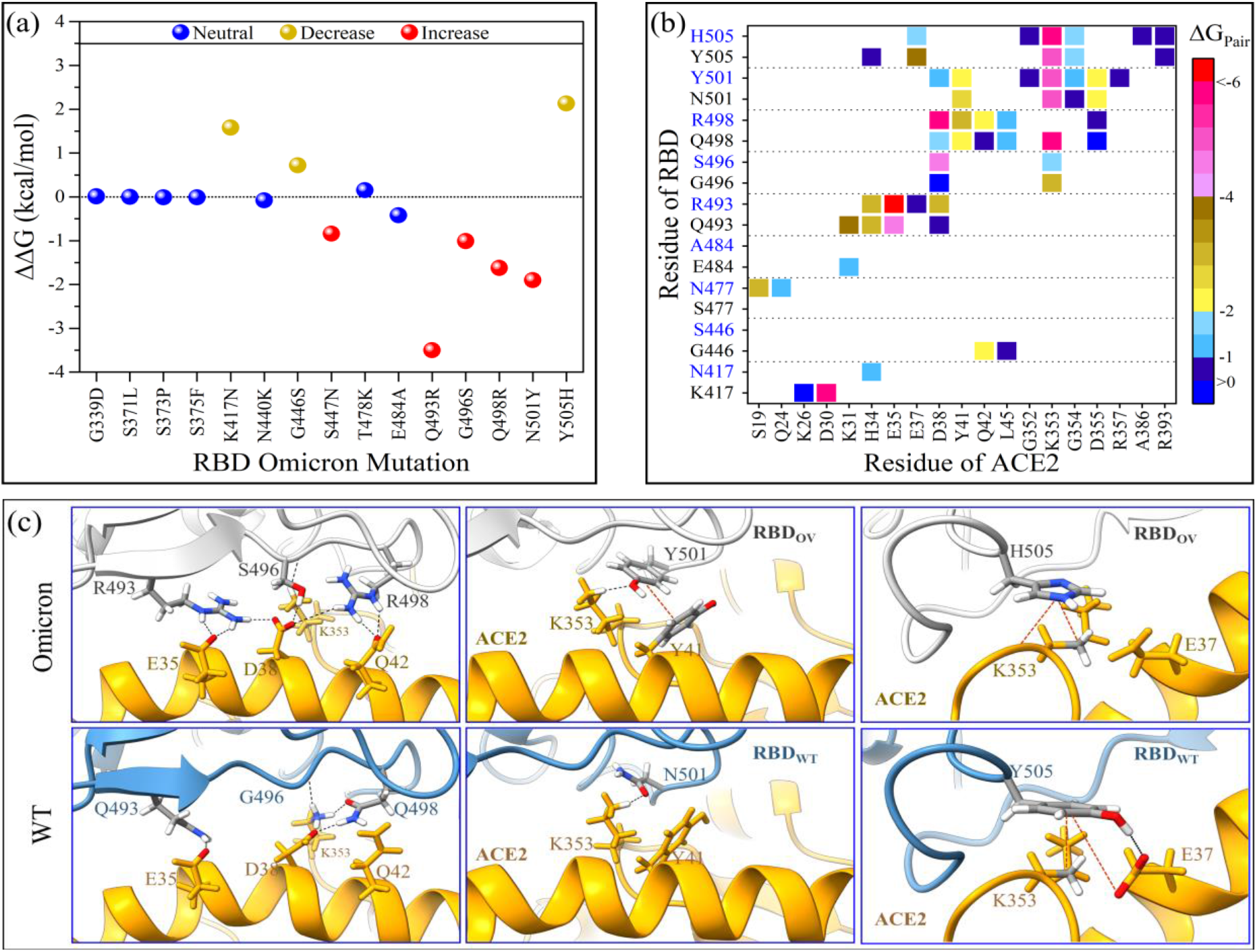
BFE decompositions of RBD-ACE2 complex in OV and WT. (a) Change in per-residue BFE decomposition (ΔΔG = ΔG_OV_-ΔG_WT_). The 15 OV mutations are classified based on their ACE2 binding properties as *neutral, increased* and *decreased* binding. (b) Pairwise BFE decomposition for only the mutated OV’s residues (blue letters in Y-axis) that form pairs with ACE2, compared with their WT (black). (c) The details of the interactions for the five important mutations in OV (Q493R, G496S, Q493R, N501Y, and Y505H) and compared with their WT. The black dash lines represent possible hydrogen bonds or salt-bridges while the red dash lines represent hydrophobic interactions.

#### 3.2.1 Mutations with no significant impact on the RBD-ACE2 association

Although mutations outside of RBM and away from the interface, such as G339D, S371L, S373P, and S375F, do not affect the RBD-ACE2 binding, they may still give rise to other biological consequences. For instance, substituting neutral G339 with highly negatively charged D339 changes the surface charge distribution of RBD (**Figure 4**), which may impair the binding between RBD and antibody. Indeed, a recent study found that this mutation has a slightly higher escape fraction for Sotrovimab, a human neutralizing monoclonal antibody (mAb).^33^ Aside from that, changing G339 to D339 results in a longer sidechain, which probably varies the local intramolecular interactions, particularly with the N343 glycosylation site.^12^ We did not include glycans in our simulation since earlier work suggested that the RBD of the S-protein had far less glycans than the S-protein itself and did not interact directly with ACE2.^34^ Mutating the polar residue (S) to the hydrophobic residues, L at 371, P at 373, and F at 375, forms a unique cluster that changes the biochemical properties of this RBD region in ways not previously observed in any other strains. This again allows OV to escape from the class 4 antibodies and some other antibodies from class 1, 2, and 3.^11,12^ Surprisingly, our MD simulations revealed that S371L, S373P, and S375F mutations are more flexible and induce a conformational change in RBD, suggesting that they have a higher chance to evade antibody recognition.

**Figure 4.**
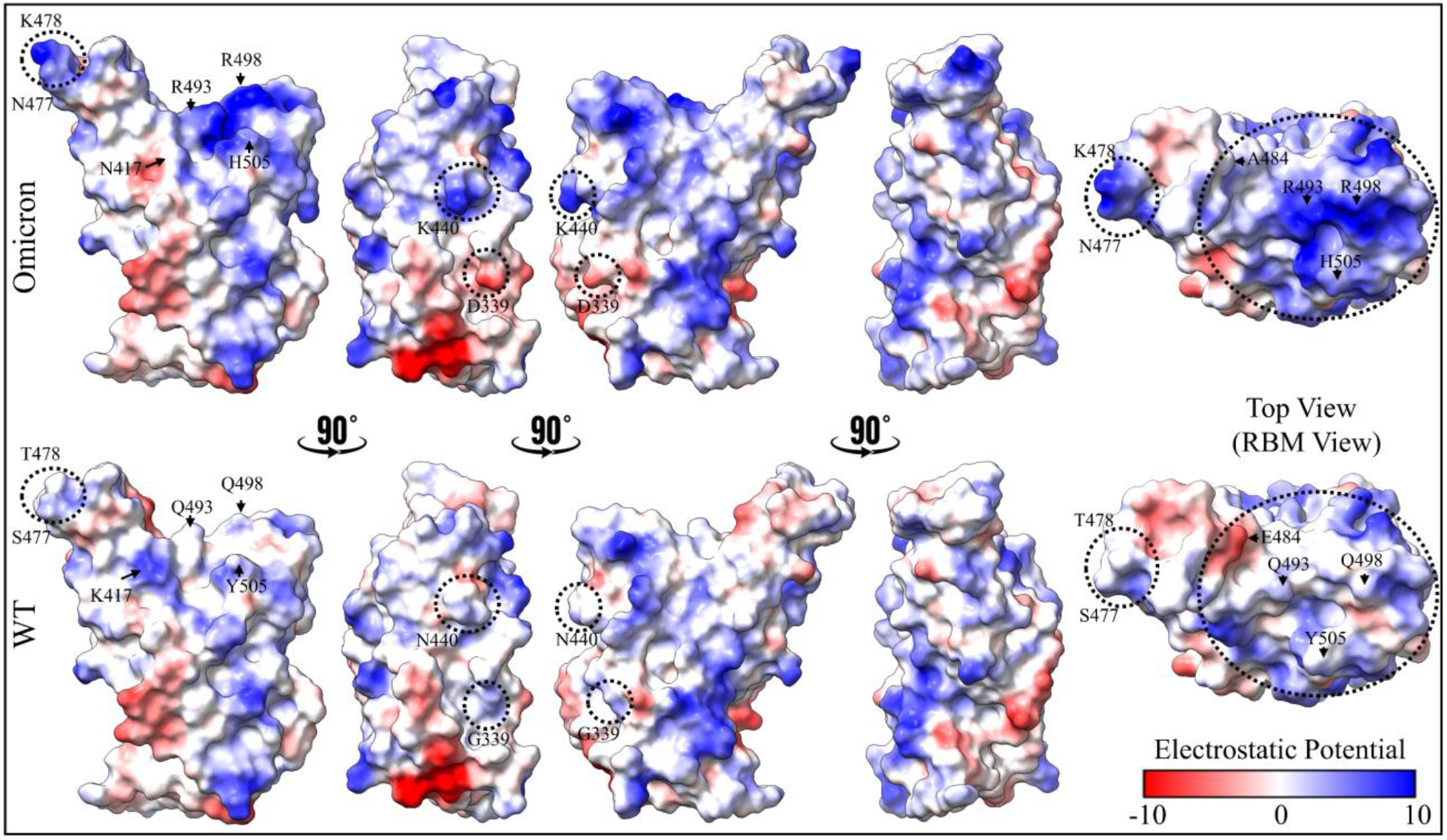
Comparison of electrostatic potential surface of RBD in mutated Omicron (top) and WT (bottom). From left to right, the first four shapes are of the RBD front view, rotated sequentially by 90°, while the fifth one represents the RBD top view. Important residues that changed their electrostatic potential surfaces are labeled. Negative-potential residues are shown in red, near-neutral residues in white, and positive-potential residues in blue.

Our findings based on BFE decompositions show that the E484A, T478K, and N440K mutations would exhibit the same pattern of neutral binding (**Figure 3**). Interestingly, E484K mutation has also been observed in Beta and Gamma VOC, and another variant of interest (VOI), and it has been identified as an *immunodominant spike protein residue*. Therefore, E484A in OV is expected to greatly reduce the susceptibility of many mAbs, which is fully consistent with the recent studies.^11,12^ In our MD simulations, we observed that the E484A mutation eliminates the weak E484:K31 in WT, but it reduces the destabilization that is stemming from possible electrostatic repulsion between E484 of WT and E35 of ACE2 when changing to A484. So, this mutation has no impact on BFE, unlike the E484K mutation in Beta, which increases the interaction marginally.^17^ T478K has also been seen in the Delta variant, and has no direct influence on the OV RBD-ACE2 interface network.^35^ Finally, the N440K mutation has been linked to an increase in antibody neutralization resistance for some antibodies.^11,36^ Importantly, the exchange of amino acids at these sites (440, 478, and 484) contributes to the emergence of a different electrostatic surface of RBD, which may play some role in attractive electrostatic interactions with the negatively charged surface of ACE2 (**Figure 4**). In this context, *ab initio* quantum methodologies offer a more accurate description of the partial charge distributions for these charged residues.^17,37–42^

#### 3.2.2 Mutations with a direct effect on RBD-ACE2 binding

Similar to the Beta variant, the K417N mutation reduces the OV RBD-ACE2 binding due to the loss of the strong ionic pair with D30 on ACE2.^17^ Certain mAbs, such as Etesevimab and Casirivimab, have been shown to be affected by K417N.^4^ Similarly, the G446S and Y505H reduce the binding, but we suspect that this is because of the impact of other mutations like N501Y rather than the mutations themselves. In our early study, we observed that the N501Y in the Alpha and Beta VOCs significantly enhances the interactions with ACE2, but it eliminates the hydrogen bond (HB) between the G446 and Q42 of ACE2 and reduces the strength of the Y505:E37 pair (**Figure 3(c)**).^17^ The same is true in OV, even though the residues at 446 and 505 site are now S and H, and as a result, their overall contributions to BFE in OV are smaller than in WT.

On the other hand, S477N, Q493R, G496S, Q498R and N501Y mutations confer additional strength to the ACE2 binding, while N477 (OV) forms new HBs with S19 on ACE2. Interestingly, our findings indicate that the main source of increased binding is due to the formation of two new strong salt bridges between R493 and R498 of OV and E35 and D38 on ACE2 with pair interaction strengths of −12.9 and −6.3 kcal/mol, respectively, which was not observed in the WT (**Figure 3(b) and (c)**). R493 (OV) can also form a salt-bridge with D38 with pair interaction strength of −2.9 kcal/mol, but it also makes a weak unfavorable pair with K353 and loses the pair with K31 that exist in the WT. R498 (OV) retains the pairings with Y41 and Q42, but they are stronger than Q498 (WT). However, it loses the pair with K353 because of the N501Y mutation.^17^ Unlike G496 of WT, the aliphatic hydroxyl group of S496 (OV) also forms new HBs with D38 on ACE2 with pair interaction strength of −4.1 kcal/mol and maintains the HB with K353 (**Figure 3(b) and (c)**). Just like in Alpha and Beta VOCs, Y501 forms more pairings than N501 of WT (**Figure 3(b) and (c)**).^17^ The number of pairs between Y501 of OV and ACE2 residues is the same as in Alpha and Beta, but their interaction strengths change, particularly the Y501:D38 pair (−0.45 of OV vs −4 kcal/mol of Alpha or Beta^17^).

In conclusion, comprehensive MD simulations have been performed to investigate the effects of OV RBD mutations on ACE2 human cell receptor binding. Our results shed light on the critical roles of the new OV mutations in the conformational changes of RBD (S371L, S373P, and S375F) and changes in its electrostatic potential surface (N440K, T478K, E484A, Q493R, Q498R, and Y505H). As a result, the overall electrostatic attraction between the positively charged OV RBD and the negatively charged ACE2 receptor is *twice as strong* as in the WT, leading to increased OV contagiousness as well as to the escape from neutralizing antibodies. Our analysis also shows that the loss of some interactions caused by K417N, G446S, and Y505H is completely compensated by the formation of new pairs from S477N, Q493, G496S, Q498R, and N501Y resulting in an overall stronger binding between OV’s RBD-ACE2.

## Supporting information

SI-BioRxiv

## ASSOCIATED CONTENT

### Supporting Information

Additional figures are provided in the Supporting Information.

## AUTHOR INFORMATION

### Author Contributions

BJ and WC conceived the project. BJ performed the calculations and made most of the figures. BJ and WC drafted the paper with inputs from PA and RP. All authors participated in the discussion and interpretation of the results. All authors edited and proofread the final manuscript.

### Notes

The authors declare no competing financial interests.

## ACKNOWLEDGMENT

This research used the resources of the National Energy Research Scientific Computing Center supported by DOE under Contract No. DE-AC03-76SF00098 and the Research Computing Support Services (RCSS) of the University of Missouri System. This project is funded by the National Science Foundation of USA: RAPID DMR/CMMT-2028803. RP acknowledges funding from the Key project #12034019 of the National Natural Science Foundation of China.

