## Supplementary material for "Binding Interactions between RBD of Spike-Protein and Human ACE2 in Omicron variant": SI-BioRxiv

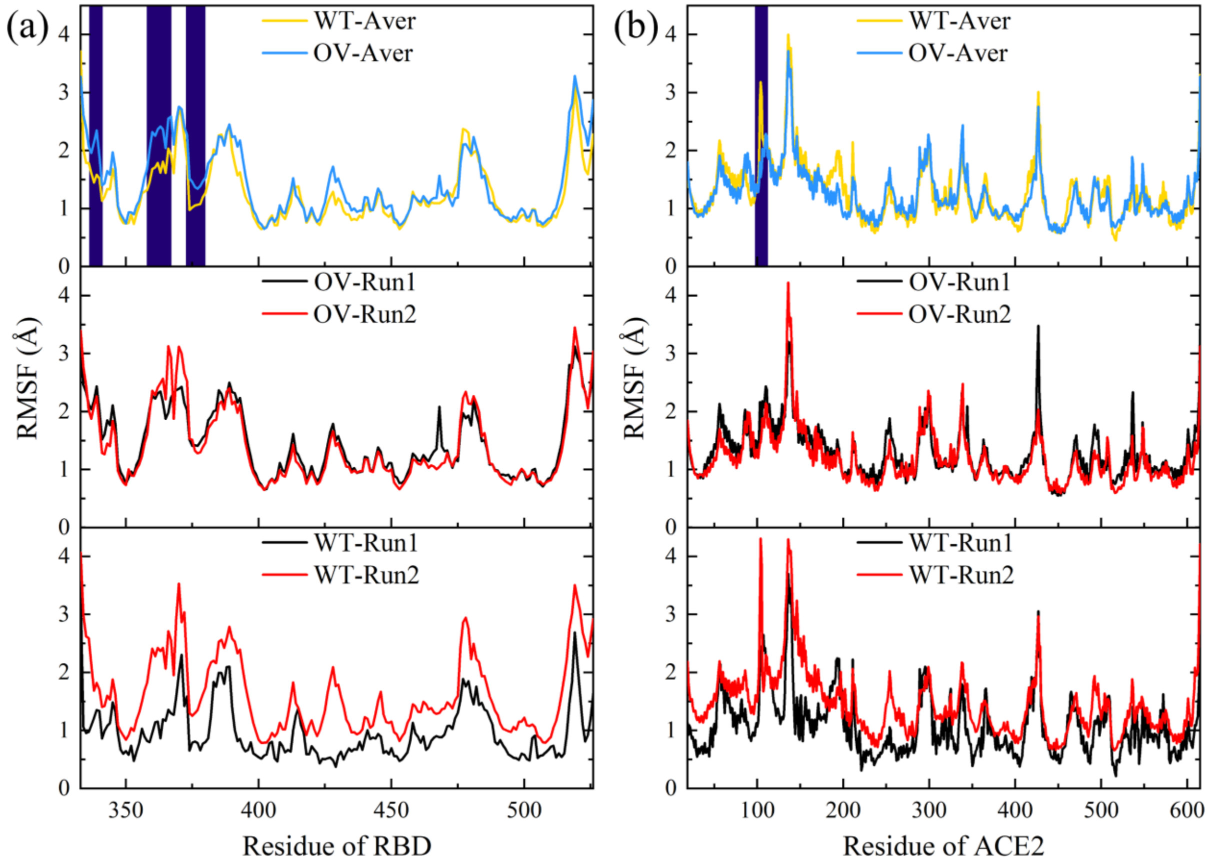


**Figure S1.** Comparison of root mean square fluctuation (RMSF) of the Omicron variant (OV) and wild-type (WT) RBD-ACE2 complexes in both MD runs (a) for RBD residues and (b) for ACE2 residues. The upper panel is for the averaged RMSF from both runs of OV vs WT. The significant RMSF differences between them are highlighted in navy. The middle panel is for OV in both runs and the bottom panel is for WT.

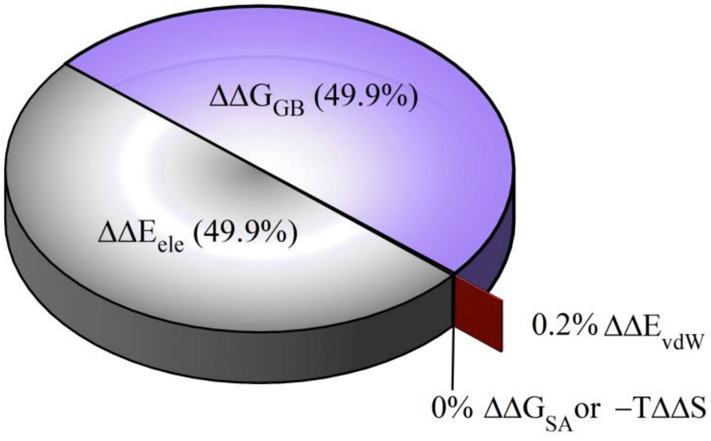


**Figure S2.** Electrostatic interaction is the main source of the difference between OV and WT.

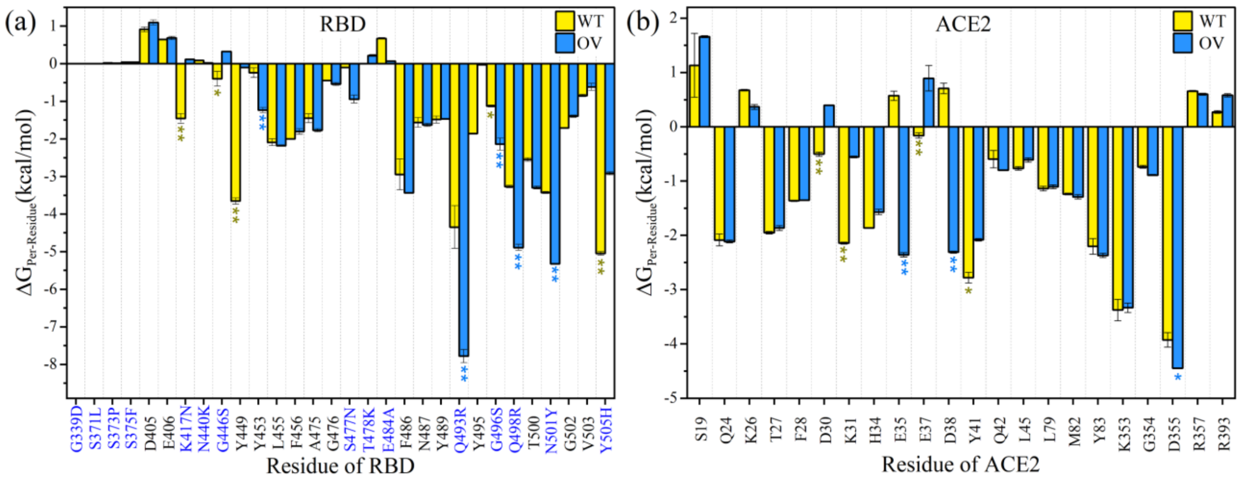


**Figure S3.** Per residue energy decomposition comparison of the interacting AAs for OV vs WT RBD-ACE2 complex. (a) For RBD residues and (b) for ACE2 residues. The significant difference is labeled with one asterisk, and the more significant differences are labeled with double asterisks.
